# Vascular topology is acutely impacted by experimental febrile status epilepticus

**DOI:** 10.1101/2022.01.10.475698

**Authors:** Arjang Salehi, Sirus Salari, Jennifer Daglian, Kevin Chen, Tallie Z. Baram, Andre Obenaus

## Abstract

Febrile status epilepticus (FSE) is an important risk factor for temporal lobe epilepsy and early identification is vital. In a rat model of FSE, we identified an acute novel MRI signal in the basolateral amygdala (BLA) at 2 hours post FSE that predicted epilepsy in adulthood. This signal remains incompletely understood and hypothesized that it might derive from changes to vascular topology. Experimental FSE was induced in rat pups and compared to normothermic littermate controls. We examined cerebral vascular topology at 2 hours, using a novel vessel painting and analysis protocol. Blood vessel density of the cortical vasculature was significantly reduced in FSE rats, and this effect was lateralized, as reported for the MRI signal. The middle cerebral artery (MCA) exhibited abnormal topology in FSE pups but not in controls. In the BLA, significant vessel junction reductions and decreased vessel diameter were observed, together with a strong trend for reduced vessel length. In summary, FSE results in acute vascular topological changes in the cortex and BLA that may underlie the acute MRI signal that predicts progression to future epilepsy. The altered vasculature may be amenable to intervention treatments to potentially reduce the probability of progression to epilepsy following FSE.

## Introduction

Febrile seizures are the most common types of seizure in human infants and young children^1^. Although most febrile seizures are generally short and have no long-term adverse effects, seizures lasting more than 30 minutes are categorized as febrile status epilepticus (FSE) and are important risk factors for developing temporal lobe epilepsy (TLE) in adulthood^2, 3^. FSE is a significant risk factor for epilepsy later in life^4^, and many adults with TLE report a history of prior prolonged febrile seizures during childhood^5, 6^. Therefore, a greater understanding of the process by which FSE in infants and children could lead to epilepsy, and especially identifying reliable markers to predict for an individual child whether he/she is at a high risk of developing TLE, is important.

A rodent model of FSE has been established and used to characterize the consequences of experimental FSE^7^. This paradigm uses immature rats at a brain development age equivalent to that of human infants and young children and induces FSE by hyperthermia that involves endogenous fever mediators^8, 9^. Like some reports in children, 30-40% of rats develop TLE in adulthood^9^. Looking for a predictive marker for epileptogenesis that may be applied in the clinic, we recently identified a novel MRI signal (reduced T2/T2* signal) in the basolateral amygdala (BLA) and hippocampus 2 hours following resolution of FSE. The signal, which was confined to one side (lateralized) was observed in ∼40% of rat pups^10^, and was highly predictive of which rats developed epilepsy in adulthood^10, 11^. The observed reduction in T2 relaxation times was a result of paramagnetic susceptibility effects, and these correlated strongly with increased deoxygenated hemoglobin. This MRI signal, denoting locally augmented deoxygenated hemoglobin might derive from increased oxygen utilization by the brain tissue or reduced blood supply acutely after FSE. The latter may be a result of abnormalities in the blood vessels, including their vasoconstriction. Because the signal was no longer present at 24-48 hours, it was considered unlikely that persistent tissue or blood vessel injury was a cause.

Thus, the underlying motivation of the current study was to identify putative mechanisms contributing to the acute reduced T2/T2* signal immediately post FSE^10^. Given that one mechanism for the altered T2/T2* signaling could be a mismatch between metabolic demand and vascular delivery, we set out to examine the vascular topology and angioarchitecture immediately after the cessation of SE in postnatal day 10 rat pups. We utilized our recently optimized vessel painting approach^12^ that directly labels the lumen of arteries and veins and thereby facilitates whole brain examination of the vascular network. We hypothesized that the vascular lumen would be reduced in vulnerable regions such as the BLA. Our experiments identified region specific alterations in vascular topology and lateralization within 2 hrs after FSE cessation, which may contribute to altered cerebral perfusion and potentially set the stage for future transition to epilepsy.

## Materials and Methods

### Animals

All animal procedures were approved by the University of California Irvine animal care committee and performed according to NIH guidelines. Sprague Dawley female dams (Envigo, Livermore, California) were maintained in quiet facilities under controlled temperatures and light-dark cycles FSE pups and their littermate normothermic controls were generated as described^7^. For technical reasons, several cohorts were used, and each cohort consisted of rat pups interleaved from different litters to minimize batch/litter effects. Within 90 minutes following resolution of FSE cessation, rat pups were removed from the dam, and all were sacrificed at the 2-hour time point. All research was conducted in accordance to ARRIVE guidelines.

### Experimental febrile status epilepticus induction

Experimental febrile status epilepticus (FSE) was induced as previously described ^7, 11^ and pups were randomly assigned to the control and FSE groups. Briefly, Sprague Dawley male and female rat pups at postnatal day (P) 10 or 11 were placed in pairs into a 3.0 litter glass container lined with absorbent paper. All pups were healthy and weighted 19.4 – 24.1 grams prior to starting FSE. Hyperthermia was induced by a continuous stream of warm air to induce behavioral seizures. Seizure behaviors progress over the hyperthermia period consisting of arrest of hyperthermia-induced hyperkinesis, chewing automatisms, and forelimb clonus. Hyperthermia (38.5-40.5°C) was maintained for 60 minutes. Core temperatures were measured at baseline, seizure onset, and every 2 minutes during hyperthermia. Following FSE, rats cooled on a chilled metal plate until their core temperature was reduced to the normal range of age. Pups were allowed to recover on a euthermic pad for 15 minutes before returning to their home cage. All pups subjected to hyperthermia experienced FSE and there were no FSE-related deaths. For normothermic controls, pups are removed from the cage for 60 minutes (to control for potential stress) and their core temperatures kept within normal range of age (34°-36C). The present study comprised of total of n=18 normothermic and n=14 FSE pups.

### Vessel painting

Vessel painting (VP) was performed as previously described ^12^. Rat pups were anesthetized with ketamine (90mg/kg) and xylazine (10mg/kg). The chest cavity was opened and Dil solution (0.45mg/ml in PBS with 4% dextrose) was manually injected into the left ventricle. Rat pups were then perfused with 10ml of phosphate buffered saline (PBS) and 20ml of 4% paraformaldehyde (PFA) at a flow rate of 7.8ml/min using a peristaltic pup. After fixation, brains were extracted from the cranium and the dura mater was carefully removed. Brains were post fixed in 4% PFA for 24 hours and stored in PBS until microscopic imaging. Criteria for including and excluding VP pups were established a priori. Successful VP was defined as uniform pink staining of the surface of the brain hemisphere and clear staining of anterior, middle, and posterior cerebral arteries as assessed by wide-field fluorescence microscopy. In the current study 11/14 FSE and 12/18 control pups had successful unilateral or bilateral staining of the hemispheres.

### Wide field fluorescent microscopy

Acquisition of wide-field cortical (axial) images were obtained by positioning the brain between two Superfrost Plus microscope slides (Fisher Scientific, Pittsburg, PA) and gently flattening the dorsal surface of the brain until a compression ring was visible on the outer edge. This allowed clear visualization of the middle cerebral artery trunk and its branching vessels. A BZ-X710 Keyence microscope (Keyence Corp XX) was used to acquire wide-field cortical images with the following settings: 2x magnification, 1 mm depth of field (25.2 µm pitch, 40 slices), and 22.5mm x 20.7mm grid size. Images were processed for maximum intensity projections using BZ-X800 Analyzer software (Version 1.1.1.8).

### Confocal microscopy

High resolution Images of the middle cerebral artery and its branching vessels from the right hemisphere were acquired with a Zeiss LSM 700 NLO laser scanning confocal microscope (Jena, Germany). Images were obtained using 10x magnification, 190 µm depth of field (10 µm pitch, 19 slices), 3.2mm x 3.2mm grid size, and Texas red filter (wavelength absorbance: 549 nm, emission: 565 nm). Images of the penetrating vessels in somatosensory cortex were acquired 1mm from the midline and obtained using a 10x magnification, 230 µm depth of field (10 µm pitch, 23 slices), and 1.9mm x 1.9mm grid size and 20x magnification and 130 µm depth of field (0.75 µm pitch, 174 slices). Photomicrographs were processed for maximum intensity projections (MIP) using Zeiss Zen software (Zeiss Zen 2010 version 1.0, Jena, Germany).

### Neurotrace Staining for BLA and hippocampal localization

The basal and lateral nuclei (BLA) are found within the anterior-posterior range from - 1.40 to -3.80mm^13^. After axial cortical acquisition and analysis was completed, brains were placed in a neonatal rat brain matrix (Zivic Instruments, BSRNA001-1) and sliced at bregma -2.40mm. A second slice was made at bregma-4.40mm to create a 2mm coronal slice. To verify the anatomical location and to delineate the boundaries of the BLA and hippocampus, the slices were stained with Neurotrace (ThermoFisher Scientific, N21480). Briefly, slices were rinsed once with PBS and then with PBS with 0.1% Triton-X for 10 minutes. Slices were incubated with Neurotrace (1/100 dilution, 500 µl total volume) for 20 minutes. Slices were rinsed with PBS with 0.1% Triton-X for 10 minutes followed by 2 rinses with PBS. Slices were left in PBS until imaging. Coronal images were obtained by positioning the slice between a Superfront plus microscope and Fisherbrand microscope cover glass (ThermoFisher Scientific, 24×60-1.5) and gently flattening the slice. BZ-X710 Keyence microscope was used to acquire images of the somatosensory cortex, hippocampus, and basolateral amygdala using 10x magnification, 243 mm depth of field (2.5 µm pitch, 97 images), and 3.5mm x 2.6mm grid size.

### Classical vessel analysis

Quantitative analysis was performed as previously described ^12^. Briefly, the raw fluorescent images were processed for black balance and haze reduction using BZ-X800 Analyzer software (version 1.1.1.8, Blur size: 20, Brightness: 20, Reduction rate: 1). In Fiji software, regions of interest (ROIs) were drawn around the right and left hemisphere on the cortical images. On coronal images the ROIs were drawn on Neurotrace images (see Immunohistochemistry below) and then overlayed onto the processed fluorescent images. The external capsule and amygdala capsule was used to outline with boundaries of the BLA^14^. Dorsal hippocampi ROIs started from the cingulum and extended to the lateral ventricles. Cortical ROIs started 1mm from the midline and outlined the brain surface and corpus callosum. ROIs were imported to Angiotool software (version 0.6a) and analysis was performed on each ROI with the following settings: vessel diameter was 10 for cortical and coronal images and vessel intensity was 15 to 255 for cortical images and 17 to 255 for coronal images. Analysis results included vessel density (percent vessels/mm^2^ * 100), junctional branch points (number of branches/mm^2^), average vessel length (average length of vessel segments), and total vessel length (total length of vessel segments). Classical vascular analysis generated a colorized AngioTool image that displays vessels in red and branch points in blue (Fig. 1).

**Figure 1.**
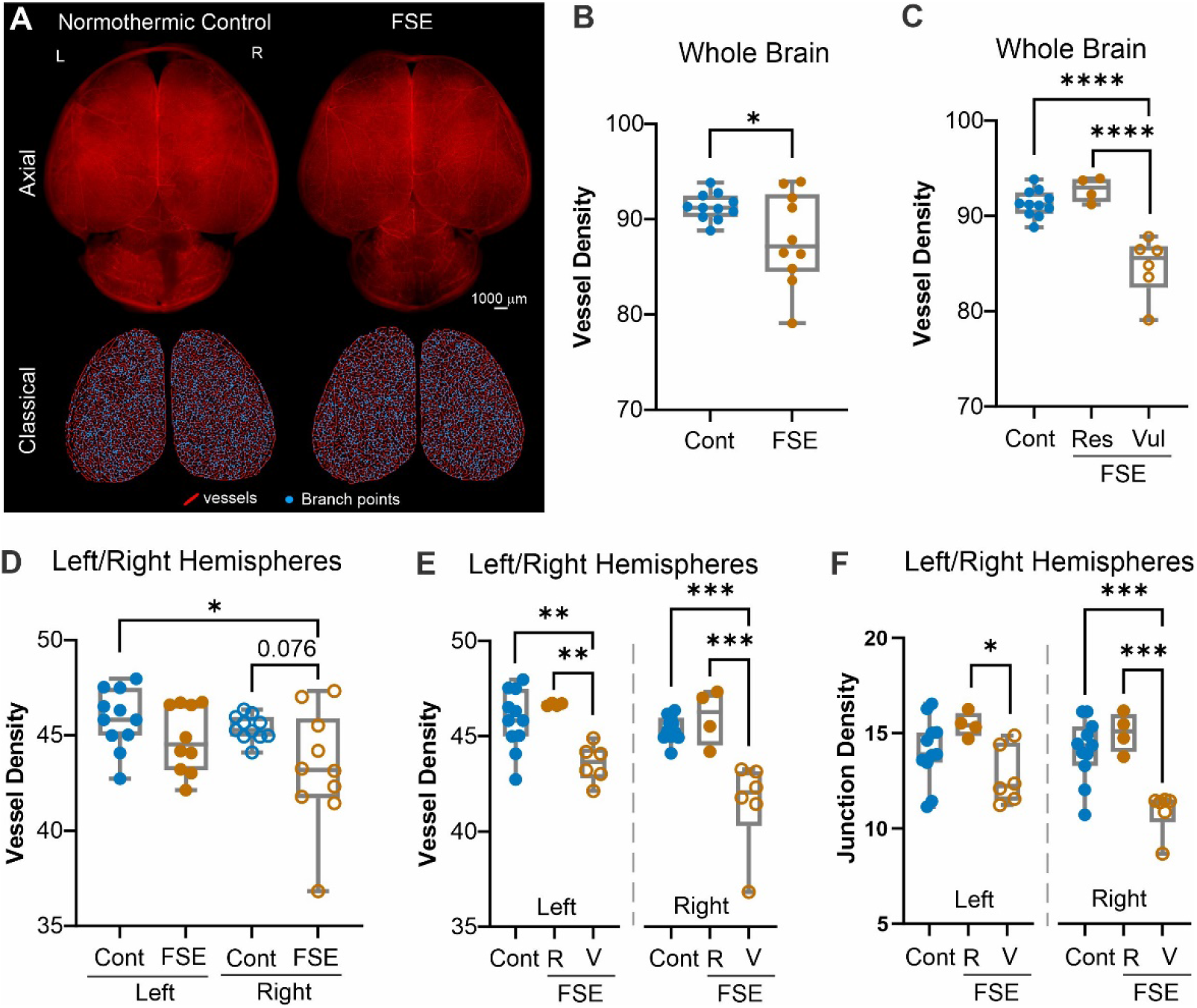
Whole brain and hemispheric vessel features. A) Vessel painting of normothermic control and FSE rat pups were undertaken at 2hrs after the onset of status epilepticus. Uniform vascular labeling occurred in both groups as shown in axial images (top panel). Vascular features were extracted from whole brain and right/left hemispheres for vessel length (red lines) and branch points (junctions, blue dots). B) Whole brain axial cortical vessel density was significantly different between groups (t-test p=0.039). However, the FSE group appeared to have two distinct groups. We have previously reported on individual vulnerability following FSE. C) Segregating the FSE pups into either low or high vessel density exhibited significant differences (one-way ANOVA, ** p<0.0001) between the vulnerable and resistant vessel density group and controls (Tukey’s post hoc, p<0.0001) and the high vessel density group (*** p<0.0001). D) When right/left hemispheres were individually examined for vessel density trending reductions (p=0.076) were observed in the right hemisphere. E) Significant differences in vessel density were found between resilient (R) and vulnerable (V) FSE rat pups, with the right side having larger decrements. (** p<0.001, *** p<0.0001) F) Vascular junction density followed similar patterns as vessel density, where the right hemisphere in vulnerable FSE rat pups appeared to have significant reductions compared to controls and FSE resilient pups. (* p<0.05, *** p<0.0001).

In our global hemispheric analyses vascular features were compared from all pups. In those cases where there was only unilateral data, group data were used to impute the missing data. Due to the unilateral or asymmetric nature of FSE-related MRI changes in children ^15, 16^ and rodents^9, 17^, we also compared the vessel features in the hemisphere with the lower values. For comparison of right vs. left hemisphere and high vs. low vascular features only pups that had bilateral uniform staining were used. From cortical images, the hemisphere with the lower vessel density value as low whereas the opposite side was designated as high. Branch points, average vessel length, and total vessel length were then dichotomized for each brain using the vessel density designations. Data from the coronal images were obtained using the low side designation derived from the cortical images.

### Fractal analysis

Fractal analyses for vascular complexity were performed using the Fraclac plugin ^18^. ROIs were drawn on the right and/or left somatosensory cortex on the confocal images. Local fractal dimensions (LFD) analysis were performed using the box counting method that results in a distribution histogram of LFD values, where local fractal dimensions (complexity) versus frequency (vessel number) are displayed. Fractal analysis generated a colorized image encoding the degree of complexity for each ROI, with a gradient of low LFD (low complexity) in purple, medium LFD (medium complexity) in green, and high LFD (high complexity) in orange. The colors are associated with complexity of the vessels, with large penetrating vessels appearing more complex than the smaller branches and capillaries.

### Vessel diameter analysis

To examine vessel diameters, extracted ROIs were processed to create tubular structures (Tubeness filter, sigma value: 10), converted to 8 bit, and thresholded (cortex images, 15 to 255; coronal images, 17 to 255). Images were then cropped to minimize black space. DiameterJ was used to obtain the mean vessel diameter and frequency histograms ^19^. We opted to use the histogram average as the mean vessel diameter as the data fell within a Gaussian distribution and the skewness and kurtosis values were between -1 and 1. The resultant histogram displays vessel diameter (µm) vs. frequency (vessel pixels along the fiber length).

### Statistics

All measurements and analysis were performed by two investigators and without knowledge of the groups. All analyses were performed using GraphPad Prism 6.0 or 8.0 (GraphPad, San Diego, CA). Outliers were identified if they were below or above 1.5 times the interquartile range. Statistical tests are noted as performed in the text. All error bars are presented as standard error of mean (SEM). Statistical significance was noted at *p<0.05, **p<0.01, or ***p<0.001.

## Results

### Global Features of Cortical Vessels

We first examined the vascular features of the entire cerebral cortex (Fig. 1). Vessel density of the entire bi-hemispheric cortices in FSE rat pups were significantly different from control pups (t-test p=0.039) (Fig. 1B). No significant differences in branch points (p=0.128), average vessel length (p=0.467) and total vessel length (p=0.543) were observed between controls and FSE pups when combined hemispheres were examined. The distribution of vascular density in both control and FSE pups was normal (Kolmogorov-Smirnov testing) but the distribution of values in FSE pups suggested two distinct clusters (Fig. 1B). When we separated those individuals with apparently resistant vessel densities from those with decreased vascular densities (vulnerable) we observed a highly significant difference between both the control and the resistant FSE groups compared to the vulnerable group (one-way ANOVA, p<0.0001) (Fig. 1C, Supplemental Fig. 1). These results were expected, because the MRI signal, which might be a result of these vessel differences, was also found only in a subgroup of FSE rats^11^.

In both rodent models^10, 11^ and humans^15, 16^, MRI changes after FSE are uniformly lateralized. Therefore, we re-analyzed the cortical vasculature separating the right and left hemispheres (Fig. 1D-F). When vessel density in FSE pups were compared to controls we found a strong trend for decreased vessel density within the right hemisphere (one-way ANOVA, p=0.076) (Fig. 1D). Specifically, FSE pups were separated into the resistant and vulnerable groups and a significant decrease (two-way ANOVA, p=0025) in vessel density was found between the control and resistant FSE pups compared to the vulnerable FSE pups, where the vulnerable FSE group was significantly different from control and high FSE pups (Tukey’s post-hoc, p<0.0001) (Fig 1E). A significant difference in vessel junction density in the right hemisphere was detected between control and resistant FSE pups when compared to vulnerable FSE pups (two-way ANOVA, p=0.04, Tukey’s post-hoc, p<0.0001). In the left hemisphere, differences were only observed between resistant and vulnerable FSE pups (p<0.05) but not compared to controls (Fig. 1F). Significant differences between the FSE and control groups were also found for average vessel length (one-way ANOVA, p=0.005; vulnerable vs control p=0.017, vulnerable vs resistant p=0.006).

Other vascular features such as average vessel length were also significantly different between hemispheres with control and resistant FSE pups having larger vessel segments on average compared to vulnerable FSE pups (Tukey’s post-hoc, left hemisphere p<0.05, right hemisphere p<0.05). Total vessel length was not altered in any of the groups. Lacunarity (space between vessels) was significantly increased in the right hemisphere of vulnerable FSE compared to control and resistant FSE pups (Tukey’s post-hoc p<0.05) but no statistically significant differences were found on the left side. In sum, FSE elicits decrements in cortical vascular features (density, junctions, average length) compared to controls. Dichotomizing the FSE into resilient and vulnerable groups clearly showed that a subset (vulnerable) of FSE pups have dramatic vascular abnormalities at 2hrs after FSE onset compared to resistantFSE pups that exhibit vascular topology like controls.

### Hemispheric Differences in Cortical Vessels

In both humans and rodents, FSE leads to MRI signal changes that are almost uniformly lateralized or even confined to a single hemisphere. In addition, in the rodent model of FSE, the right hemisphere appeared to be more involved^17^. We plotted right/left vascular features for each control and FSE pup for comparison (Fig. 2A). Hemispheric differences were not observed in controls (p=0.977, FSE p=0.153) in vessel density (Fig. 2A). In FSE rats, there were significant differences between the two hemispheres, but they did not involve the right or left hemisphere consistently (t-test, p=0.009) (Fig. 2A). Importantly, in the more impacted hemisphere we observed highly significant decreases in vessel density in vulnerable FSE pups compared to FSE resistant and control groups (Fig. 2B). Interestingly, the vulnerable FSE pups had reduced vessel density clustered entirely on the right hemisphere unlike the resistant FSE pups which were equally distributed.

**Figure 2.**
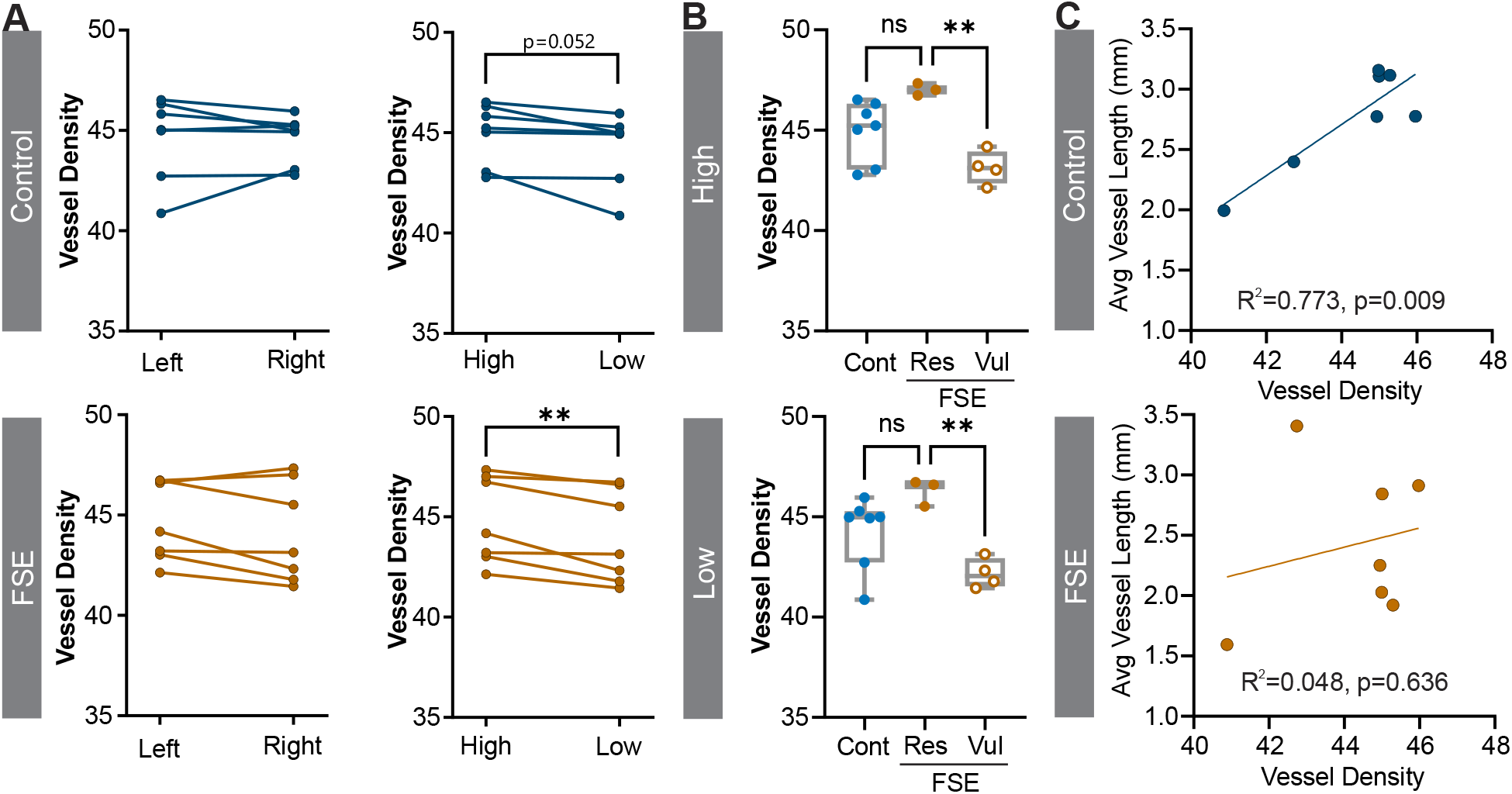
Lateralization in FSE. A) Right and left hemispheres showed no significant differences in vessel density between controls and FSE pups (left panel). When vessel densities were dichotomized into high vs. low vessel density control pups approached significance (p=0.052) but FSE pups were strongly significantly different (**p=0.009) (right panel). B) Further separation based on the vulnerable and resistant FSE pups further demonstrated highly significant reductions in vessel density in the vulnerable group compared to FSE (High **p=0.004, Low **p=0.007). C) Correlations between the low hemisphere vessel density and average vessel length were strongly correlated in control pups (R^2^=0.773, p=0.009) but was disrupted in FSE pups (R^2^=0.048, p=0.636).

Vascular junctional density was not significantly different between right and left hemispheres. When right and left hemispheres were dichotomized into high junctional density vs. low junctional density, both FSE pups and controls exhibited a significant decrease in junction density compared to the high hemisphere (6.96 ± 1.26, p=0.004 control; 6.02 ± 1.06, p=0.003 FSE). Within FSE pups there was a significant junction density difference between vulnerable and resistant pups (p=0.012) but not to controls. Thus, junctional density does not differentiate between FSE pups and controls.

The right hemisphere of every FSE pup had reduced total vessel length compared to the left side. Accordingly, total vessel length was significantly reduced in the right hemisphere of FSE pups (p=0.010) compared to the left with no significant differences in controls. In contrast, average vessel length was not significantly different between hemispheres in control or FSE pups. When hemispheric average vessel length data were sorted into high vessel length vs. low vessel length values there was a significant difference in controls (p=0.023) and in FSE pups there was a more prominent decrease in average vessel length (p=0.003).

In control pups there was a strong correlation between vessel density and average vessel length (R^2^=0.773, p=0.009). Notably, this correlation was not found in FSE pups (R^2^=0.048, p=0.636), as shown in Fig. 2C.

### Middle Cerebral Artery Characteristics

The middle cerebral artery (MCA) is one of the most prominent cortical vessels and supplies the amygdala, a primary site of MRI signal abnormalities. (Fig. 3A). We measured the diameter of the MCA branches (M2-4) in controls and FSE pups (Fig. 3A. Supplemental Fig. 2)) and no significant differences were found between groups (two-way ANOVA, p=0.846). As expected, reductions in vessel diameter were found within both control and FSE pups (p<0.0001) as we moved to increasing distal branches (Fig 3B). There were no significant MCA branch X Group effects (p=0.748). Examination of FSE pups for resilient vs. vulnerable dichotomization found no significant differences after repeated measures ANOVA testing (p=0.089). However, resilient FSE MCA vessel diameters tended to be larger than controls and vulnerable FSE vessels (Fig. 3C). Vessel diameter frequency histograms did not identify overt global changes in vessel diameters when either right, left or lower vessel diameter histograms were examined (Fig 3D).

**Figure 3.**
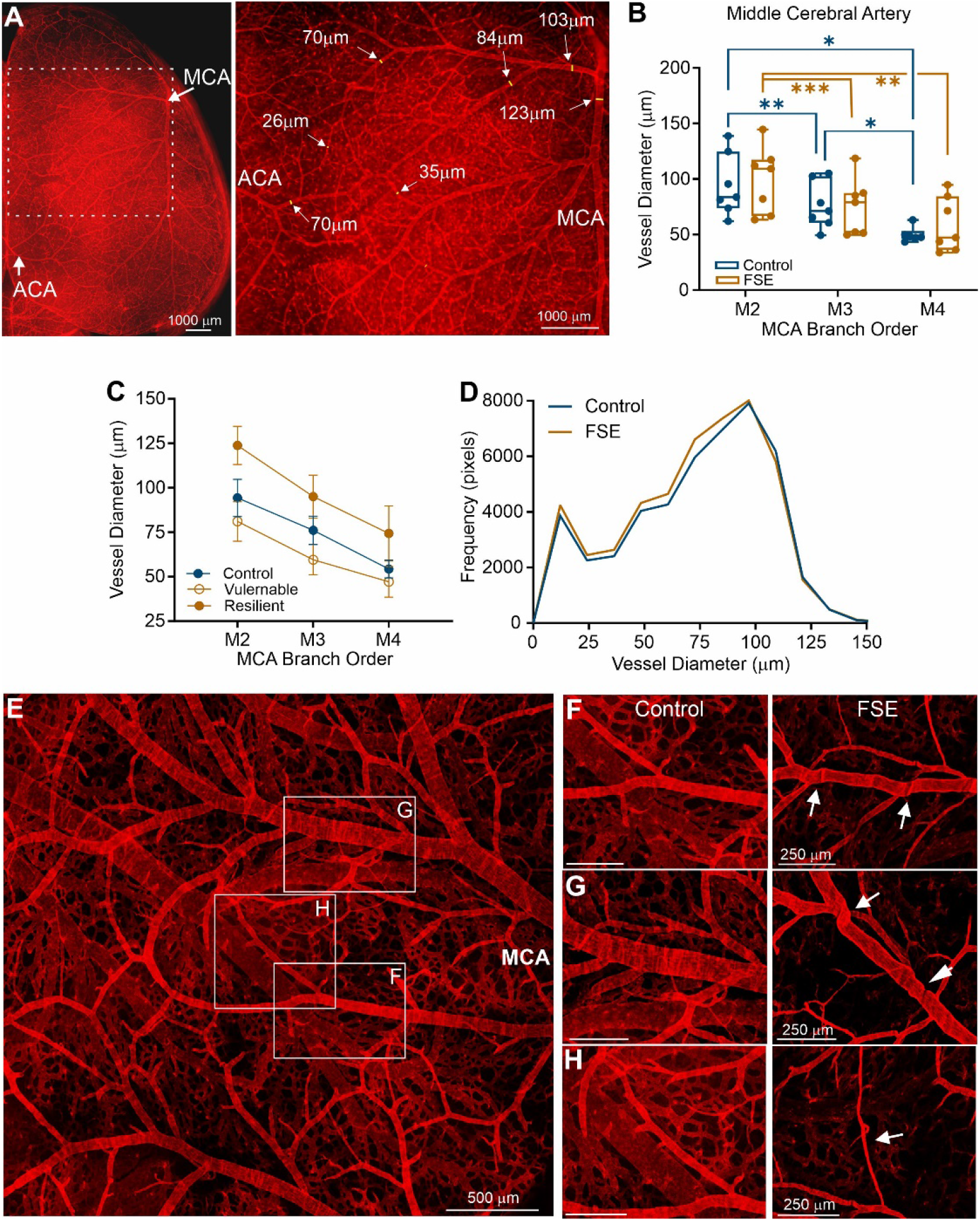
Vascular features of the middle cerebral artery (MCA). A) The most prominent vascular structure on the axial surface of the brain is the MCA. Diameter measurements were applied to confocal images from vessel painted brains. B) There were no significant differences between control and FSE pups in vessel diameters but both groups progressed to small diameters with increased branching (M2-4). C) Resilient and vulnerable FSE groups were compared and while the resilient group had elevated diameters compared to controls and vulnerable FSE pups, these findings were not significant. D) Frequency histograms of vessel diameter did not exhibit any overt differences when comparisons were made to lower hemispheric vessel diameters. E) In control pups the MCA territory exhibited a robust vascular network with normal appearing vascular elements. F-H) However, comparisons of vessels between control (left panel) and FSE (right panel) exhibited robust differences in FSE animals with vessel folding (F), tortuosity at junctions (G) and vessel thinning (H).

While quantitative MCA metrics were not overtly different between control and FSE pups, significant qualitative vascular abnormalities between the groups were visible. In control pups there was a substantial vascular network present within the MCA territory region that is comprised of a distribution of larger and small vessels (Fig. 3E). Closer examination in the controls showed normal vascular diameters, branching and vascular density (Fig. 3F-H, left panels). The vascular features in controls were in stark contrast with the features from the MCA in FSE pups (Fig. 3F-H, right panels). Many FSE vessels in the MCA territory had apparent kinks (Fig 3F, right), tortuosity at vascular bifurcations (Fig. 3G, right) and numerous thin vessel segments (Fig. 3H, right). Thus, while global features may be only moderately altered, examination of the vascular network at high magnification revealed dramatic differences in vessel angioarchitecture, and these resembled changes reported in several brain pathologies^20^.

### Vessel Characteristics of the Basolateral Amygdala

The basolateral amygdala (BLA) appeared to be uniquely sensitive to FSE and MRI signal changes in this region predicted of long-term development of epilepsy^10, 11^. A random subset of whole vessel painted brains (n=4/group) were blocked at the level of the dorsal hippocampus which included the BLA (∼ Bregma-2.40mm) (Fig. 4A,B). First, we identified group (control vs FSE) differences in the total number of vessel junctions (mixed model, p=0.025) (Fig. 4C). Vascular characteristics within the right versus left BLA were not overtly different between control and FSE pups, including vessel density (one way ANOVA p=0.30), average vessel length (p=0.237) and junctional density (p=0.255). We noted a strong trend for differences in vessel area p=0.076 and total vessel length (p=0.085). We then divided for each pup the high vs low hemisphere / BLA as above and identified a significant inter-hemispheric difference in the number of junctions in FSE but not control pups (post-hoc t-test, p=0.047) (Fig. 4D).

**Figure 4.**
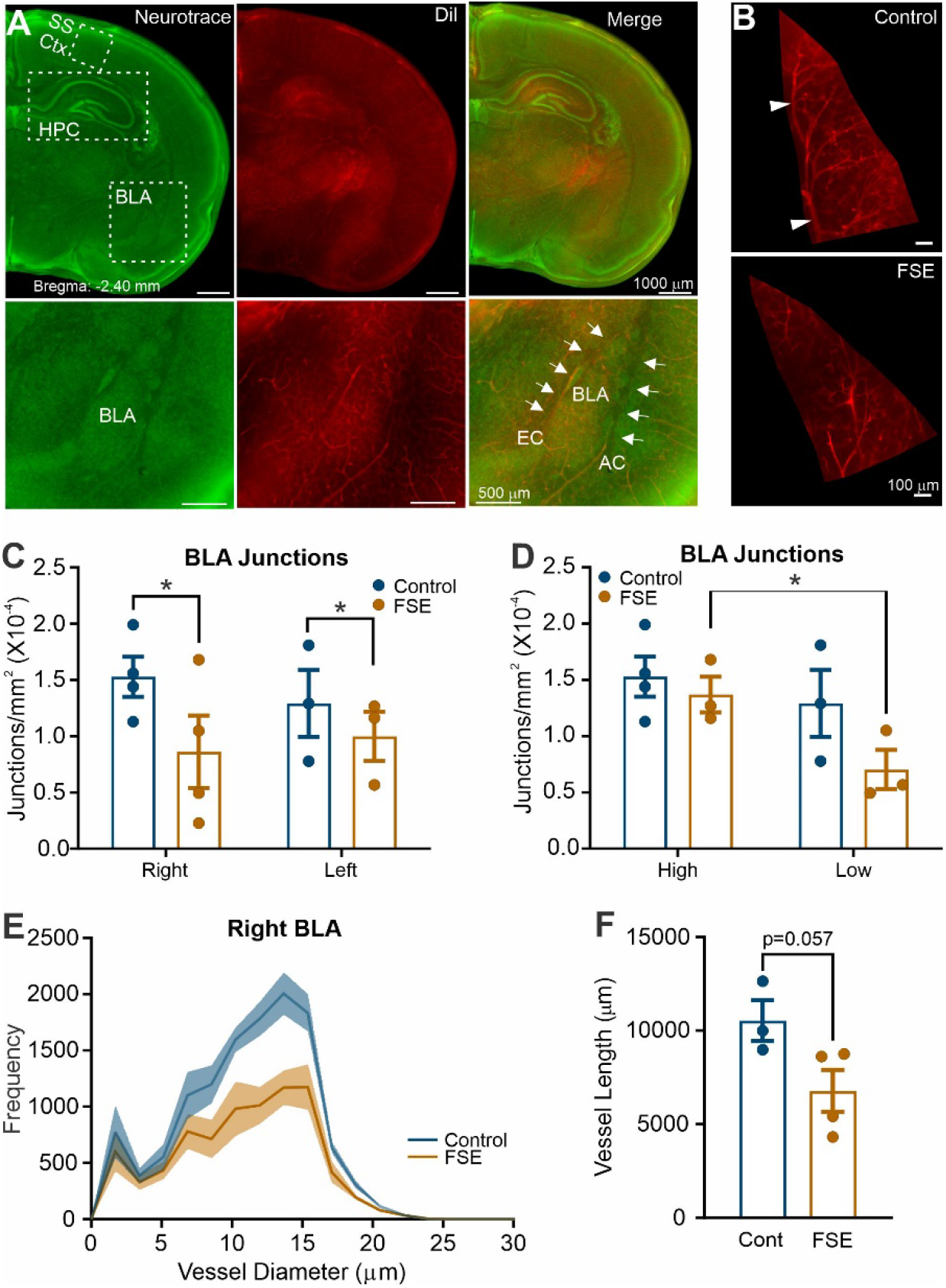
Basolateral amygdala vessel features. A) Representative coronal slices at the level of the dorsal hippocampus for somatosensory cortex (SS Ctx), hippocampus (HPC) and basolateral (BLA) vessel analyses. Left panel is sections stained for Neurotrace to assist in region localization. The middle panel is the vessel painted image from the same section and the right panel is the merged images. The bottom row of images was taken at higher magnification to show localization of the BLA. B) Representative vessel painted images from the right BLA of a control (top) and FSE (bottom). As can be seen vascular density is reduced. C) Vessel junctions were in the right and left BLA were significantly different between groups (Control vs FSE; mixed model, *p=0.025) but not differences between right and left. D) Junctions were then dichotomized to high and low with no significant differences between controls (mixed model, p=0.126), but in FSE pups the low side was significantly reduced compared to the high side (post-hoc t-test, p=0.047). E) Vessel diameters within the BLA were also quantified with a 55.5% decrease in the area under the curve (t-test, p=0.067) but only 17.7% on the left BLA (data not shown). F) Trending significant reductions in total vessel length were observed.in the right (but not left) BLA (t-test, p=0.057

Diameters of vessels in the BLA were robustly (55.5%) decreased in 10-15μm sized vessels (Fig.4E). In the right BLA area under the curve analysis found a trend for decreased vessel diameters between control and FSE curves (t-test, p=0.067) (Fig. 4E) with no significant differences in curve skewness or kurtosis. There was a trend for decrease in the frequency of vessels sized 11-20 μm (Mann-Whitney, p=0.057) in the right BLA. There were no significant differences between in vessel diameters <11 or >20 μm. In contrast with the right hemisphere, there was a non-significant 17.7% decrease in vessel diameters in the left BLA based on area under the curve.

Fractal analysis for vessel complexity did not find significant differences between control and FSE pups, although there were reduced numbers and complexity in FSE pups compared to controls (data not shown), This trend was more prominent in the right compared to the left BLA.

### Vessel Characteristics of the Hippocampus

Hippocampus is also a site of altered MRI signal after FSE ^10, 17^, and is highly involved in MRI signal changes in the human^16, 21^. Here, a trend for an interaction between groups (control vs. FSE) and right vs left hippocampus (mixed effects, p=0.054) was noted. Significant interactions were observed for average vessel length (mixed effects, p=0.006) and junction density (mixed effects, p=0.032) between control and FSE pups (Fig. 5A). Of note, vessel length was increased in left FSE hippocampus compared to control pups (post-hoc testing p=0.027), and junction density trended higher (post-hoc test p=0.079) (Supplemental Fig. 3).

**Figure 5.**
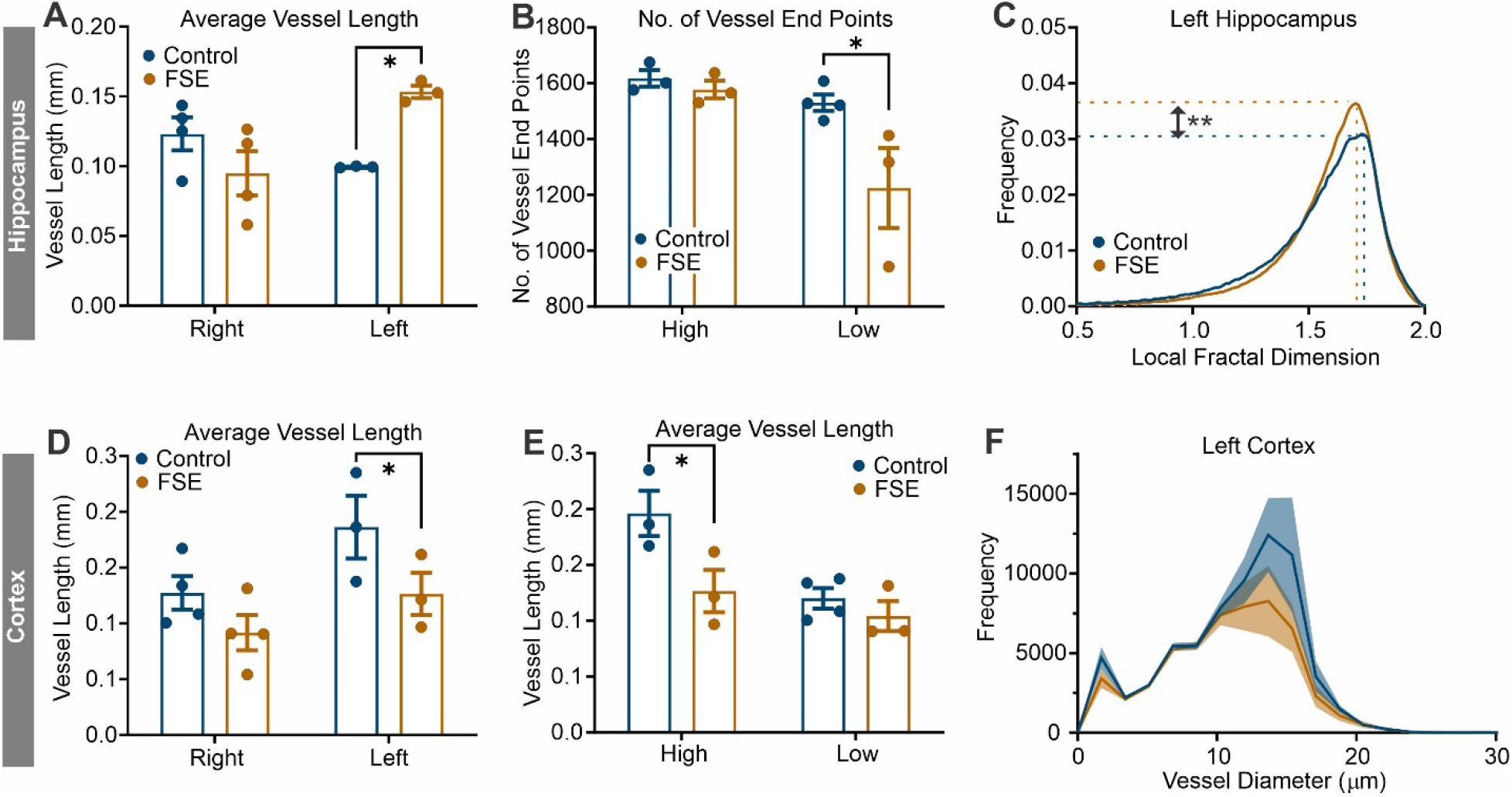
Vascular features of the hippocampus and cortex. A) Average vessel length was increased in FSE pups compared to controls in the left hippocampus (mixed effects post-hoc, p=0.027). B) The number of vessel end points were significantly reduced in FSE pups compared to controls when dichotomized into high/low groups (mixed effects post-hoc, p=0.023). C) In fractal analysis the left hippocampus of FSE pups displayed increased peak frequency (t-test, p=0.007) consistent with increased vessel numbers (see A). D) In the cortex, average vessel length was significantly differ in the left cortex between control and FSE pups (*p=0.032). E) Average vessel length was then grouped into high/low designations and there was a significant reduction in the high FSE group compared to controls (mixed effects, *p=0.012). There were no differences between FSE high and low lateralization in the cortex. F) Vessel diameter histogram for the left cortex illustrates reductions in vessel diameter in the FSE pups compared to controls; this reduction was not significant.

As in the BLA, we compared high vs. low hemispheric vessel features. Vessel density was significantly lateralized (mixed effects, p=0.0002), with both controls and FSE pups having similar high/low values (data not shown). Similar inter-hemispheric differences in junction density (mixed effects, p=0.0003), vessel area (mixed effects, p=0.015), total vessel length (mixed effects, p=0.023) and average vessel length (mixed effects, p=0.002) were observed. Only the number of vessel end points (terminations) was significantly reduced in FSE compared to controls (mixed effects, p=0.018; post-hoc p=0.023) (Fig. 5B). There were no significant changes in vessel diameters in either the right or left hippocampus of FSE pups compared to controls.

Fractal analysis was performed to buttress the classical neuroanatomical approach (Fig. 5C). Maximum frequency was elevated in the left hemisphere (t-test, p=0.007) but not the right hemisphere (t-test, p=0.707) of FSE compared with controls. The maximum frequency is considered a measure of vessel number^12^. Similarly in the left hemisphere skewness and kurtosis were significantly different from controls (t-tests, p=0.007, p=0.015, respectively) with a trend for changes in the right hemisphere (skewness p=0.065, kurtosis p=0.071). These fractal changes within the hippocampus are in accord with and confirm the increased classical vascular features.

### Somatosensory Cortex

The somatosensory cortical region immediately above the hippocampus was also assessed for vascular modifications. There were no significant changes in vessel area, total number of junctions, vessel junction density, total vessel length between control and FSE pups nor between right and left hemispheres. However, there was a significant difference in average vessel length between right and left hemispheres (mixed effects, p=0.034) and between FSE and control pups (mixed effects, p=0.032) and no significant interaction between group and hemisphere (mixed effects, p=0.544) (Fig. 5D).

When hemispheres were dichotomized into high and low vessel features, total number of junctions (mixed effects, p=0.005), junction density (mixed effects, p=0.001) and average vessel length (mixed effects, p=0.012) were significantly different between groups and high/low designations. In all significant parameters the high designation was significantly different between the groups, where vascular features were significantly lower in FSE pups than controls (Fig. 5E). The number of vessel end points was higher in FSE compared to control pups, but this was not significant (mixed effects, p=0.177). The increased average vessel lengths potentially suggests that there may be increased vessel fragmentation or reduced branching in the cortex as other vascular parameters were not significantly different between controls and FSE pups.

Vessel diameter histograms showed reductions in vessel diameters with distance from vessel origin in both the left and right cortex but were not significantly different between controls and FSE pups (Fig. 5F). Similarly, no other vessel diameter parameters were significantly altered. Accordingly, fractal analyses found no significant differences in any measure within the somatosensory cortex

## Discussion

These studies report on novel and acute changes in vascular topology after experimental febrile status epilepticus. Notably, these changes provide a potential mechanism for the acute MRI signals observed in a subset of rats after experimental FSE, changes that predict the future development of epilepsy

Using a hyperthermic model of FSE, we report acute changes in the vascular topology. The major findings in FSE pups are: 1) global reductions in cortical vascular density wherein a subset demonstrated highly significant reductions (FSE vulnerable). 2) Global changes were lateralized, i.e. one hemisphere was more affected than the other. 3) The MCA territory vessel topology exhibited tortuosity, thinning (string-like) and abnormal kinks at the vessel bifurcations. 4) The BLA exhibited reductions in junctional density related to reduced vascular length. 5) The hippocampus had lateralized increases that contrasted with decreased average vessel length in the somatosensory cortex. In summary, the novel predominate finding was that there were reductions in vascular elements (density, junctions etc) acutely after FSE. These acute modifications may have implications in the development of subsequent epilepsy.

Altered vascular topology has been reported both acutely and long-term in adult models of status epilepticus ^22^. To our knowledge there are no hyperacute (hours post SE) studies examining the vascular morphology nor studies examining the vascular consequences of FSE. Thus, Nodode-Ekane in a pilocarpine model of status epilepticus (SE) found reduced hippocampal vessel density, vessel length accompanied by increased blood brain barrier (BBB) leak at 2 days post SE using immunohistochemical staining^23^. Other models of SE, including chemical induction of SE in adult rodents resulted in vessel degeneration^23, 24^ and leakage of the BBB^23, 25-27^, which are primarily evident in the hippocampus, amygdala, thalamus, and parietal cortex^23, 25, 27^. Fabene and colleagues found that within 2 hrs after adult rat pilocarpine induced SE onset that cortical vessels exhibited flattened morphology but that had increased vascular diameters (∼20%)^28^. Interestingly, in deeper cortical layers vascular diameter was decreased by 46% and related to decreased cerebral blood volumes. The authors also noted cell death in the upper cortical layers but not in the lower layers. Using a similar model of pilocarpine induced SE but at postnatal day (PND) 9 and 21, Marcon and colleagues mapped vessel density at 7 days post SE^29^. Using immunohistochemistry (FTIC-albumin, laminin) they found no change in hippocampal, amygdala nor cortical vessel density in PND9 rat pups compared to controls. In contrast, at 7 days post SE in PND21 rats there was a 30% increase in hippocampal vessel density but only in a subset (62%) of SE rats; the other SE rats were similar to controls. Similar elevations in vessel density were found in cortex (38%) and in amygdala (30%)^29^. In combination with other evidence the authors suggest that these increases in vessel density 7 days after SE are the result of ongoing angiogenesis. Similar angiogenic mechanisms were proposed by Ndkode-Ekane and colleagues after pilocarpine induced SE in adult rats using immunohistochemical methods^23^. At 2 days post SE there was decreased vessel density and length within the hippocampus (CA1/CA3) that only marginally improved over the 60-day experimental period. However, there was a robust increase in BrdU-labeled endothelial cells at 2, 4 and 14 days after SE, that correlated with increasing vessel length and suggested an angiogenic process^23^. In brain trauma we have demonstrated a strong neo-vasculogenesis mechanism that appears to be independent of vascular endothelial growth factor (VEGF) ^30^. The role of VEGF in epilepsy has recently been reviewed^31^. Our own data demonstrate that cortical vessel density (Fig. 1) is already significantly decreased within 2 hrs after FSE, with similar reductions in the BLA (Fig. 4) and increases in the hippocampus (Fig. 5). While it is difficult to compare across models and time points, SE results in perturbations of vessel topology. In our model the cortical disruptions in vessel density were also accompanied by disruption of the BBB^32^, and associated with decreases in vessel junctions (branching) confirming that changes in vascular angioarchitecture may contribute to ongoing and future pathophysiology of FSE.

A second novel aspect of the altered vascular network was the apparent lateralization within the cortex of the vascular decrements. There were no significant differences between right/left hemispheres vessel density in controls (0.02%) or FSE (1.67%) pups. In our previous MRI studies, we observed that there was lateralization of the T2 biomarker signal^10, 11^. We used a similar strategy in our vascular assessments (high vs. low values) to lateralize these findings. When lateralization was performed FSE pups had a 2% reduction in vessel density (high vs. low) that was significant whilst controls had a 1.6% change that was trending. While no studies exist on vascular lateralization there is some support for lateralization of blood flow in seizures, with increases during ictal discharges and decreased blood flow after seizure cessation that was asymmetric between hemispheres^33^. We recently reported similar asymmetric differences in MRI T2 signal and cerebral blood flow in a preliminary study of FSE patients that underwent MRI evaluations within 6 hrs of SE onset^34^. Cerebral blood flow has been also shown to lateralize with pathology in newborns with focal seizures^35^. Thus, while there is scant literature on vascular lateralization, future studies should examine if alterations are asymmetric and as such may have the potential for diagnostic interpretation.

In human sclerotic hippocampi there was a 60-70% reduction in the number of microvessels that were also accompanied by a wide variety of vascular abnormalities^36^. The vasculature of the sclerotic CA1 had a rough appearance with spine-like protrusions and abnormal tubular structures that were associated with severe neuronal loss and gliosis. In two mouse models of epilepsy, microvascular vasospasms were increased relative to control mice with increased mural cell numbers and vessel constrictions^37^. The authors went on to demonstrate that capillary vasospasms increased 400% within 80 sec prior to seizure onset, suggesting that vascular constriction may lead or certainly contribute to seizure onset. These vessel spasms then lead to local hypoxia with subsequent cell death^37^. In our model of FSE there is no overt neuropathology, but within the MCA territory we observed tortuous vessels, increased vascular “kinking” or abnormal folding of the vessels adjacent to vessel branches (Fig. 3G-H). While we cellular basis for these abnormalities is uncertain, it is tempting to speculate that capillary pericytes that are associated with vessel branches may contribute to the altered morphology^38^. Similar modified pericyte structures in tissues excised from human temporal lobe epilepsy patients were associated with tortuous and abnormal vessels, frequently at vessel junctions^39^. Thin or string-like vessels, like those we observed in the cortex of FSE pups (Fig 3H), have been associated with a variety of disease states^20, 40^. These abnormal structures have not been rigorously studied in SE and specifically in FSE but likely represent ongoing vascular pathology that may impact long-term patient outcomes.

An interesting aspect of our study is that vessel painting allows brain-wide vessel staining. Thus, in addition to cortical assessments, we can probe specific regions of interest, such as the hippocampus, BLA and somatosensory cortex above the hippocampus (Figs 4, 5). We found generalized decreases in vascular features in the BLA and cortex but increases in the hippocampus. Increased vascular densities were found in the cortex and amygdala, days and weeks after adult rat pilocarpine and kainite-induced SE^29, 41^. Within the hippocampus the overwhelming evidence is for increased microvascular density days to weeks after induced adult SE (see Table 1 in Marcon^22^ for a concise summary). In our study we found increased vessel length in the hippocampus of FSE pups at 2 hrs after the cessation of SE that was mirrored by our fractal analyses (Fig. 5C). Functionally, cerebral blood flow and volume was not altered in the hippocampus at 2 days post pilocarpine SE but was increased at 14 days when vessel density increased^23^. In human temporal lobe epilepsy bilateral decreases in cerebral blood flow were found in unilateral epilepsy which was unrelated to neuronal loss^42^. Given the paucity of studies on regional vessel density, use of different models and time points it is difficult to craft a picture of how the vasculature responds to SE. Our own data provide a glimpse into the acute vascular response to FSE but clearly additional studies are warranted.

Several limitations of the current study are noted. Firstly, temporal monitoring of vascular topology over chronic and latent epileptogenic epochs would provide insights into the dynamic nature of the vascular response to FSE. Secondly, functional measures of blood flow using laser doppler or MRI perfusion imaging would have also enhanced the interpretation of our results. Finally, immunostaining for pericyte markers would have provided insights into the abnormal morphology observed in the MCA as well as other markers, such as smooth muscle.

The current studies employed a preclinical model of FSE, in which an acute MRI signal presaged the development of TLE-like limbic epilepsy. Recently, the same signal has been observed in infant with FSE^34^, but it is not yet clear if the same signal predicts epilepsy in children with FSE. This possibility, now under study^34^, provides strong impetus for identifying the underpinnings of the predictive signal that associates with augmented deoxygenated hemoglobin. The abnormal vessel topology found here, including tortuosity and reduced diameters and vessel density, provide a plausible and important mechanism for the MRI observation, potentially leading to strategies for intervention.

## Authors Contribution Statement

Concept and Design: AS, TZB, AO

Data Acquisition: AS, JD, KC

Data Analysis: AS, SS, AO

Data Interpretation: AS, TZB, AO

Manuscript Drafts and Revisions: AS, TZB, AO

## Acknowledgements

These studies were supported by an NIH Training grant from the National Institute of Neurological Disorders and Stroke (T32 NS-45540). This study was made possible in part through access to the Optical Biology Core Facility of the Developmental Biology Center, a shared resource supported by the Cancer Center Support Grant (CA-62203) and Center for Complex Biological Systems Support Grant (GM-076516) at the University of California Irvine.

## Notes

### Competing Interest Statement

The authors have declared no competing interest.

