## Supplementary material for "Vascular topology is acutely impacted by experimental febrile status epilepticus": Supplimental Figures

**Supplemental Figures**

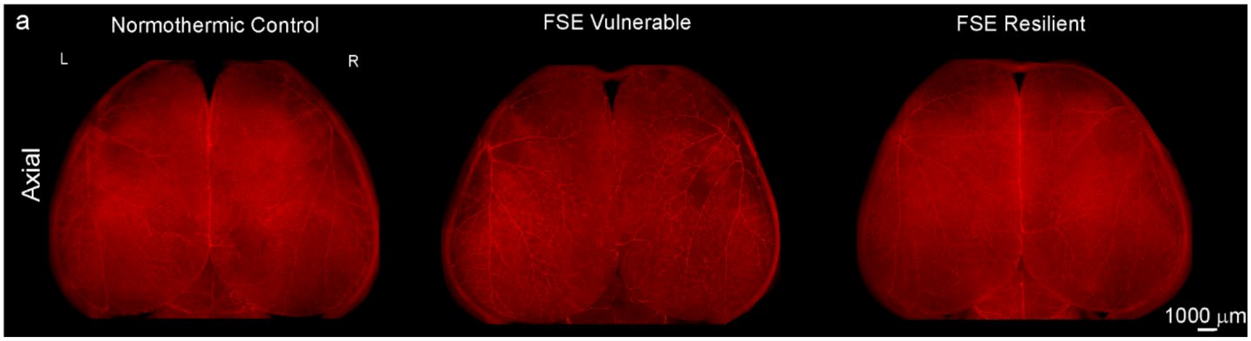


**Supplemental Figure 1. Exemplar vessel painted brains from control and FSE pups.** Visually it is apparent that the vulnerable FSE pup (center image) has reduced vascular density compared the resistant FSE pup (right image). The FSE resistant vessel painted brain resembles the control brain (left image).

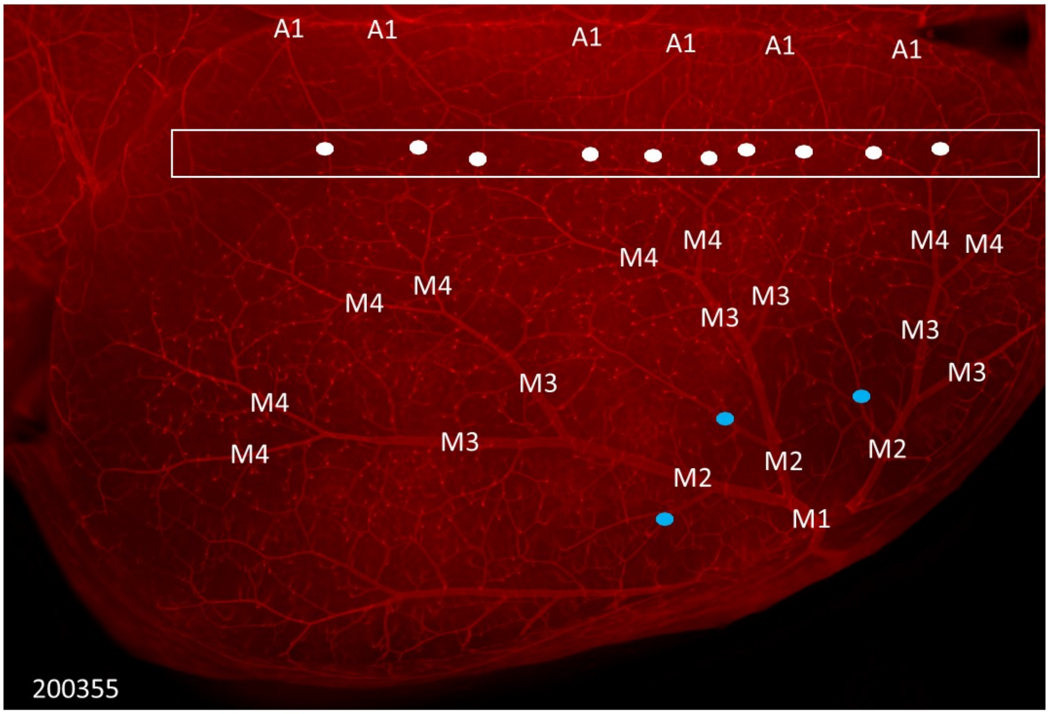


**Supplemental Figure 2: Middle cerebral artery (MCA) branches for vessel diameter analyses.** We labeled all the branches (M2-4) of the MCA on the hemispheric surface and measured the diameters of the vessels within each branch. A1= anterior cerebral artery (ACA). White dots = region of ACA and MCA anastomosis. Blue dots = regions of MCA anastomosis that were not included in our diameter measurements.

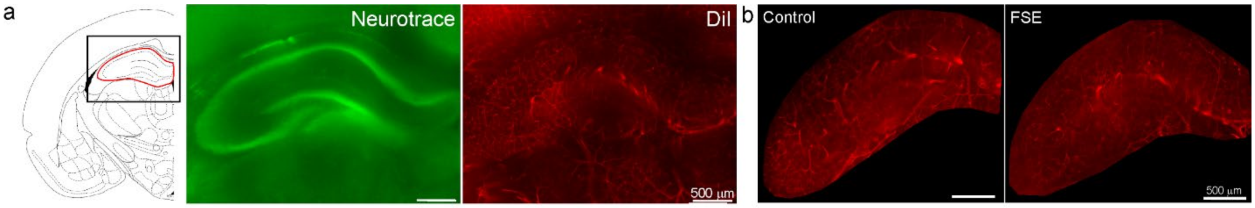


**Supplemental Figure 3: Hippocampal vessel painting.** A) The hippocampus was identified on from the Neurotrace images, and a region of interest was drawn that was then duplicated to the vessel painted (DiI) hippocampus. B) The hippocampal regions were then excised from the whole brain vessel painted images for subsequent classical and fractal vascular analyses.
